# Sex-Specific Regional Brain Morphometric Correlates of Neighborhood Socioeconomic Disadvantage in Clinical Neuroimaging

**DOI:** 10.64898/2026.08.10.743987

**Authors:** Ethan H. Willbrand, Luis A. Vazquez, Michael B. Fromandi, Paloma C. Frautschi, Troy B. Powell, John-Paul J. Yu

**Affiliations:** Medical Scientist Training Program, University of Wisconsin School of Medicine and Public Health, Madison, WI, United States; Department of Radiology, University of Wisconsin School of Medicine and Public Health, Madison, WI, United States; Neuroscience Training Program, University of Wisconsin School of Medicine and Public Health, Madison, WI, United States; Department of Biostatistics and Medical Informatics, University of Wisconsin School of Medicine and Public Health, Madison, WI, USA; Department of Psychiatry, University of Wisconsin School of Medicine and Public Health, Madison, WI, United States; Department of Biomedical Engineering, College of Engineering, University of Wisconsin-Madison, Madison, WI, United States

## Abstract

**BACKGROUND AND PURPOSE:** Neighborhood-level socioeconomic disadvantage is associated with adverse brain morphometry, yet whether these associations differ by biological sex remain opaque. Here, we investigated sex-specific associations between the area deprivation index and brain morphometry derived from routine clinical MRI in a real-world clinical population.

**MATERIALS AND METHODS:** Intracranial volume-normalized regional brain volumes were extracted from T1-weighted MRI examinations performed in 2,863 consecutive clinical patients (median age 54 years [IQR 38–68]; 61.2% female) at a single academic medical center and associated community partners using an automated atlas-based segmentation pipeline. Exploratory factor analysis was applied to 131 regional brain volumes to identify latent neuroanatomical morphometric networks. Sex-stratified linear regression models examined associations between area deprivation index national percentile rank and each factor score, adjusting for age, with correction for multiple comparisons.

**RESULTS:** Factor analysis identified five neuroanatomical morphometric networks: cerebellar (ML1), frontal/executive (ML2), subcortical-ventricular (ML3), medial temporal/limbic (ML4), and posterior cortical/visual (ML5). In male patients (*n* = 1,112), linear regressions revealed that greater neighborhood-level socioeconomic disadvantage was significantly associated with lower factor scores on the cerebellar (β = −0.006, 95% CI [−0.009, −0.003], *P* < .001), medial temporal/limbic (β = −0.004, 95% CI [−0.007, −0.001], *P* = .01), and frontal/executive (β = −0.004, 95% CI [−0.007, −0.0004], *P* = .04) networks. No significant associations were observed in female patients (all *P*s ≥ .61).

**CONCLUSIONS:** In a real-world clinical population, neighborhood-level socioeconomic disadvantage was associated with lower regional brain volumes across cerebellar, frontal/executive, and medial temporal/limbic neuroanatomical morphometric networks in male but not female patients. These findings suggest that the neuroanatomical correlates of neighborhood disadvantage may be sex-specific, and that sex-stratified analyses may be necessary to fully characterize the relationship between the social exposome and brain morphometry in clinical neuroimaging research.

## INTRODUCTION

The life-course social exposome, the cumulative environmental and socioeconomic conditions in which individuals live throughout their lives, has emerged as a critical determinant of health.^1^ Among the multitude of factors comprising the social exposome, neighborhood-level socioeconomic disadvantage represents a particularly salient measure that is able to simultaneously encode structural inequities in income, education, employment, and housing that extend beyond individual-level risk factors.^2^ Neighborhood-level socioeconomic disadvantage has been consistently linked to adverse health outcomes, including increased cardiovascular disease burden, psychiatric morbidity, and accelerated cognitive decline.^3,4^ Concomitantly, recent neuroimaging studies have begun to demonstrate that such exposures are associated with alterations in brain structure, primarily presenting as reduced global brain volume, and in some cases, reduced regional brain volumes (e.g., the hippocampus and prefrontal cortex).^5–11^

However, much of this work has been derived from enriched research cohorts or disease-specific samples, limiting generalizability to real-world clinical settings.^5–10^ In a recent epidemiology study leveraging a large, consecutive real-world clinical sample, our group demonstrated that residence in socioeconomically disadvantaged neighborhoods was associated with higher brain age gap and lower total brain volume, as well as amplified associations between white matter hyperintensity burden and regional brain volumes.^11^ These findings suggest that neighborhood context is not merely a background demographic variable, but rather a measurable correlate of brain morphometry detectable in routine clinical neuroimaging.

Considering these advances, a persistent gap in knowledge is whether the impact of neighborhood disadvantage on the brain differs by biological sex. Extensive prior work has established the role of biological sex in multiple domains directly relevant to brain health, including early neurodevelopment, aging, vascular risk, and stress responsivity, as well as the prevalence and presentation of neuropsychiatric and neurodegenerative disorders.^12–15^ Recent evidence further suggests that neighborhood disadvantage may differentially influence brain structure across sexes, with sex-specific associations observed in cortical regions implicated in stress, emotion regulation, and pain processing in samples enriched for disease (e.g., irritable bowel syndrome^10^ and Alzheimer’s disease^6^). These differences raise the possibility that the biological embedding of socioeconomic disadvantage may not be uniform across sexes but instead may manifest in sex-specific patterns of brain morphometry. Crucially, failure to account for sex-specific effects may obscure meaningful associations in neuroimaging studies and limit the interpretability of findings in clinical practice. Given that biological sex has been shown to systematically influence brain structure and disease risk, omission of sex-specific analyses may lead to biased estimates or the masking of true associations.^12,14,16^ As neuroimaging-derived measures of brain morphometry are increasingly applied to assess brain health across the lifespan, understanding how these metrics vary across biological and social dimensions is essential for accurate risk stratification.^17–19^

Motivated by these findings, in the present study, we extend prior work by investigating whether the association between neighborhood-level socioeconomic disadvantage, measured using the area deprivation index (ADI),^2^ and brain morphometry differs between male and female patients in a large, consecutive, real-world clinical population.^11^ Specifically, we applied exploratory factor analysis to intracranial volume-normalized regional brain volumes derived from routine clinical T1-weighted MRI to identify latent neuroanatomical morphometric networks spanning cerebellar, frontal/executive, subcortical-ventricular, medial temporal/limbic, and posterior cortical/visual networks. We then examined sex-stratified associations between ADI national percentile rank and each neuroanatomical morphometric factor score, adjusting for age, in a cohort of 2,863 patients. These analyses may provide insight into how structural socioeconomic inequities are differentially reflected in brain morphometry across sexes, with implications for the integration of social and biological context into neuroimaging research and clinical interpretation.

## MATERIALS AND METHODS

### Study Sample

Participant data were retrospectively obtained from consecutive clinical MRI examinations performed in both inpatient and outpatient settings at a single academic medical center and associated community partners between January and June 2024. The initial dataset included 35,704 individuals. Exclusion criteria were applied sequentially: patients were removed if imaging reports indicated structural or pathological abnormalities (e.g., neoplasm, infarction, demyelinating disease) or if they were younger than 18 years. Patients were also excluded if they lacked neighborhood disadvantage or morphometric data. These data were previously used by our group to assess the relationship between ADI and brain morphometry, not explicitly considering sex differences.^11^ In-depth details on the inclusion/exclusion process are provided in this prior work.^11^ The final analytic cohort consisted of 2,863 participants [median (IQR) age: 54 (38–68) years; 61.2% female; **Table 1**]. This study followed the Strengthening the Reporting of Observational Studies in Epidemiology reporting guideline.

**Table 1:** Participant demographic characteristics and neuroanatomical morphometric factor scores by sex.

| Variable | Overall<br>(*n* = 2,863) | Male<br>(*n* = 1,112) | Female<br>(*n* = 1,751) | *P*-value |
| --- | --- | --- | --- | --- |
| **Demographics** |  |  |  |  |
| Age, years — median (IQR) | 54 (38–68) | 59 (41–70) | 52 (36–67) | < .001 |
| ADI national percentile — median (IQR) | 41 (30–53) | 41 (30–55) | 41 (31–52) | .04 |
| **Neuroanatomical Morphometric Factor Scores — median (IQR)** |  |  |  |  |
| ML1: Cerebellar | 0.014<br>(–0.613–<br>0.627) | 0.004<br>(–0.632–<br>0.635) | 0.021<br>(–0.585–<br>0.615) | .32 |
| ML2: Frontal/Executive | 0.053<br>(–0.562–<br>0.604) | 0.088<br>(–0.622–<br>0.693) | 0.023<br>(–0.534–<br>0.564) | .17 |
| ML3: Subcortical-Ventricular | 0.085<br>(–0.612–<br>0.675) | –0.085<br>(–0.755–<br>0.571) | 0.168<br>(–0.509–<br>0.721) | < .001 |
| ML4: Medial Temporal/Limbic | –0.026<br>(–0.613–<br>0.594) | –0.082<br>(–0.728–<br>0.492) | 0.021<br>(–0.544–<br>0.639) | < .001 |
| ML5: Posterior Cortical/Visual | –0.025<br>(–0.642–<br>0.643) | 0.007<br>(–0.600–<br>0.682) | –0.047<br>(–0.673–<br>0.618) | .11 |

**Note:**—ADI = area deprivation index; IQR = interquartile range; ML = maximum likelihood factor. *P*-values for age and ADI national percentile are from independent-samples *t*-tests. Factor score *P*-values are from independent-samples *t*-tests comparing male and female participants.

### Quantifying Neighborhood Disadvantage

Neighborhood-level socioeconomic disadvantage was quantified using the ADI, a composite metric derived from 17 census-based indicators encompassing income, education, employment, and housing characteristics. ADI values were obtained from 2023 American Community Survey data via the publicly available Neighborhood Atlas (https://www.neighborhoodatlas.medicine.wisc.edu/).^2^ Each participant’s most recent residential address was geocoded using ZIP+4 information and mapped to a corresponding census block group, allowing assignment of national-level ADI rankings.^2^

### MRI Processing and Morphometric Extraction

Structural MRI data were processed using a fully automated, atlas-based segmentation pipeline (cNeuro, Combinostics Oy, Tampere, Finland). T1-weighted images were nonlinearly registered to a set of reference atlases, and regional labels were propagated and combined to generate final segmentations. This multi-atlas approach has been previously validated in neurodegenerative disease cohorts for reliable volumetric quantification.^20^ Regional volumes were calculated based on voxel counts within each labeled structure and scaled by voxel dimensions. Composite measures were generated by aggregating relevant regional outputs. To account for interindividual variability in head size, all volumetric measures were normalized using intracranial volume-based scaling.^21^ Age- and sex-adjusted normative percentiles were also derived using established reference distributions.^21^ In line with prior work,^22,23^ left and right hemispheres were modeled separately for the subcortical and cortical regions to capture potential hemisphere-specific differences.

### Statistical Analyses

All statistical analyses and data visualization were implemented in R (v.4.6.0; https://www.r-project.org/).

#### Factor Analysis of Regional Brain Volumes

Factor analysis of regional brain volumes identifies latent morphometric covariance networks, that is, sets of regions whose volumes co-vary systematically across individuals, analogous to structural covariance networks described in prior neuroimaging research.^24–26^ To reduce the dimensionality of the regional morphometric data and identify latent neuroanatomical morphometric networks, exploratory factor analysis was applied to intracranial volume-normalized regional brain volumes. From the full set of atlas-derived volumetric measures, 131 regions of interest were selected for inclusion, comprising bilateral subcortical structures, cerebellar regions (including vermal lobule groups), and cortical regions of interest spanning frontal, temporal, parietal, occipital, and insular cortices (see **Supplemental Data** for the complete list of included regions). Prior to factor extraction, all variables were z-scored. The factorability of the resulting correlation matrix was assessed using the Kaiser-Meyer-Olkin (KMO) measure of sampling adequacy (overall MSA = 0.50) and Bartlett’s test of sphericity (χ²(8515) = 355,729.1, *P* < .001).^27,28^ Although the KMO value was at the lower threshold of acceptability,^27,28^ Bartlett’s test confirmed that the correlation matrix was significantly different from an identity matrix, and factor analysis proceeded given the large sample size and the known modest intercorrelations among spatially distributed brain regions inherent to whole-brain volumetric data,^24–26^ and the interpretability of the resulting solution.^29^ The number of factors to retain was determined by inspection of the scree plot in conjunction with parallel analysis (maximum likelihood estimator; 100 iterations; see **Supplemental Data** for the scree plot). Parallel analysis formally suggested retention of 30 factors; however, this recommendation is known to over-extract in large samples with many weakly correlated variables. Guided by the scree plot (**Supplemental Data**), which demonstrated a clear inflection point after the fifth factor, and by the anatomical interpretability of the resulting solution (**Fig 1**), five factors were retained. Factor extraction was performed using maximum likelihood estimation with oblimin rotation, selected *a priori* to permit correlated factors consistent with the anticipated covariance structure of regional brain morphometric data. Factor scores were computed using the regression method.

**Fig 1.**
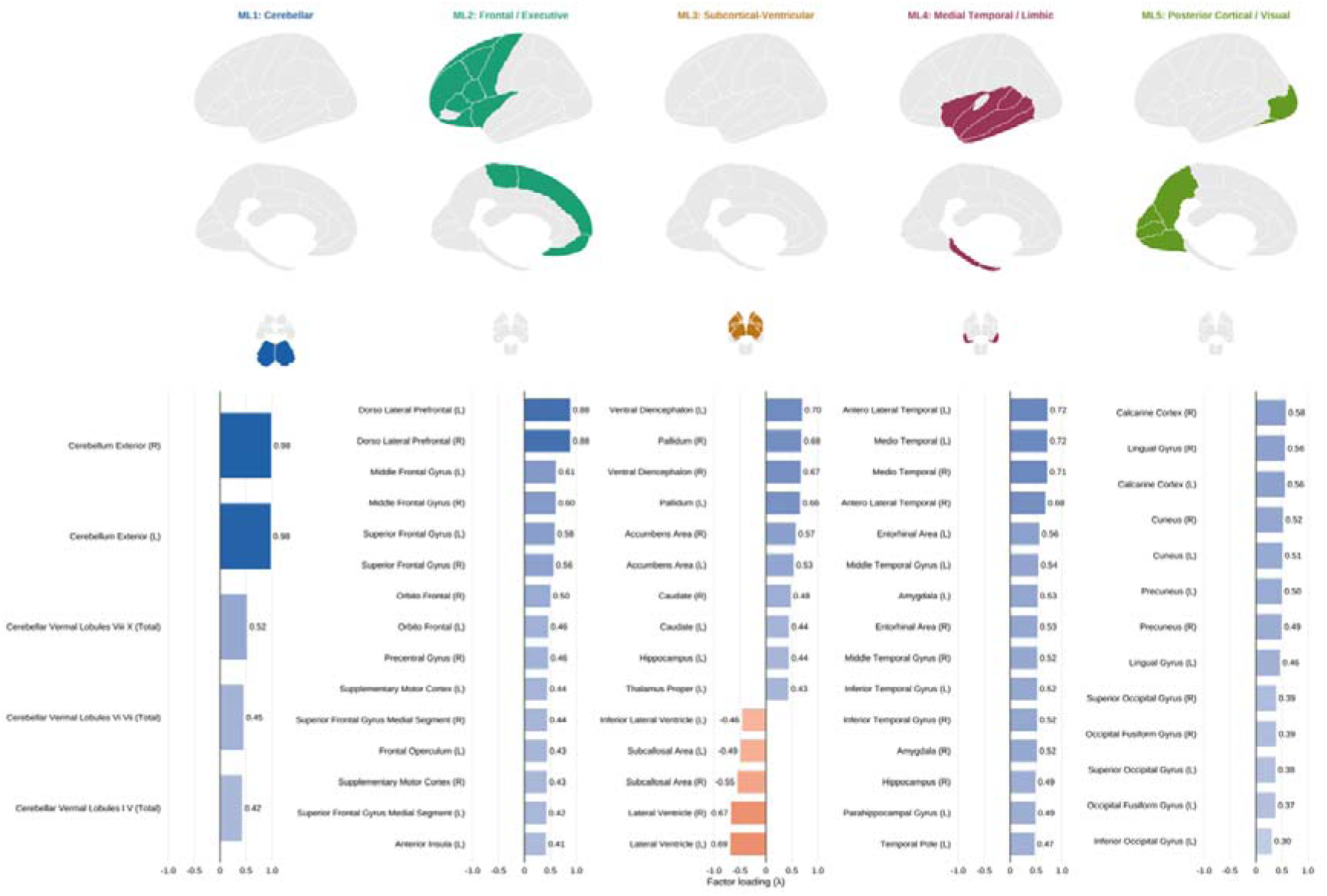
Exploratory factor analysis of regional brain volumes identifies five neuroanatomical morphometric networks. *(A)* Brain schematics are provided as a visual aid to orient readers to the approximate neuroanatomical distribution of each neuroanatomical morphometric factor. Top row: Left hemisphere lateral cortical view. Middle row: Left hemisphere medial cortical view. Bottom row: Subcortical axial view (ML1: axial slice 3; ML2–ML5: axial slice 4). Regions are colored based on approximate mapping of higher-loading regions (|λ| ≥ 0.30) between the atlas used in the automated segmentation pipeline (cNeuro, Combinostics Oy) and the Desikan-Killiany cortical atlas for cortical views (https://surfer.nmr.mgh.harvard.edu/fswiki/CorticalParcellation) and the FreeSurfer subcortical segmentation atlas for subcortical views (https://surfer.nmr.mgh.harvard.edu/fswiki/FreeSurferVersion3); refer to these atlases for complete region nomenclature. *(B)* Bar charts display the top 15 regions per factor with |λ| ≥ 0.30 (when possible), arranged by loading magnitude. Blue bars indicate positive loadings; orange bars indicate negative loadings. Complete factor loadings for all 131 regions of interest are provided in the Supplemental Data. ML = maximum likelihood factor; λ = factor loading.

#### Regression Model Implementation

To examine whether neighborhood-level socioeconomic disadvantage was associated with each latent neuroanatomical morphometric factor, and whether these associations differed by biological sex, linear regression models were implemented separately for male and female participants. For each of the five factors, ADI national percentile rank and age were entered as predictors. Scanner was not included as a covariate because all scans were acquired on a harmonized fleet of GE clinical scanners using standardized protocols across sites. To account for multiple comparisons across the five factor models within each sex, *P*-values for the ADI coefficient were adjusted using the Benjamini-Hochberg false discovery rate (FDR) procedure. Statistical significance was defined as FDR-adjusted *P* < .05.

## RESULTS

### Participant Characteristics

The final analyzed cohort comprised 2,863 participants with a median (IQR) age of 54 (38– 68) years, of whom 1,751 (61.2%) were female and 1,112 (38.8%) were male. Median (IQR) ADI national percentile rank was 41 (30–53) for the overall sample, 41 (30–55) for male participants, and 41 (31–52) for female participants; ADI national percentile rank differed significantly between sexes (t(2861) = 2.09, *P* = .04). Male participants were significantly older than female participants [median (IQR): 59 (41–70) vs. 52 (36–67) years; t(2861) = 5.32, *P* < .001]. Full demographic and neuroanatomical morphometric factor score characteristics are presented in **Table 1**.

### Factor Structure of Regional Brain Volumes

Exploratory factor analysis of the 131 intracranial volume-normalized regional brain volumes yielded five neuroanatomical morphometric networks that together captured the principal axes of morphometric covariance across the cohort. The five-factor solution was selected based on the scree plot inflection point and anatomical interpretability, as described in the **Materials and Methods**. The top regions for each factor, which helped term the five profiles described below, are shown in **Figure 1**. Factor loadings for all 131 regions are provided in the **Supplemental Data**.

The five factors exhibited distinct neuroanatomical morphometric profiles (**Fig 1**). The first factor (ML1) was defined by strong positive loadings from bilateral cerebellar hemispheres and all three vermal lobule groups, consistent with a “cerebellar neuroanatomical morphometric network” (**Fig 1**). The second factor (ML2) was characterized by high positive loadings across dorsolateral prefrontal, middle and superior frontal, orbitofrontal, precentral, and supplementary motor cortices, reflecting a “frontal/executive neuroanatomical morphometric network” (**Fig 1**). The third factor (ML3) captured a “subcortical-ventricular axis,” given positive loadings from the pallidum, ventral diencephalon, nucleus accumbens, caudate, putamen, and thalamus, and negative loadings from lateral ventricles and subcallosal area (**Fig 1**). The fourth factor (ML4) was dominated by anterolateral and mediotemporal cortices, entorhinal area, amygdala, hippocampus, parahippocampal gyrus, and temporal pole, consistent with a “medial temporal/limbic neuroanatomical morphometric network” (**Fig 1**). The fifth factor (ML5) was defined by positive loadings from calcarine cortex, lingual gyrus, cuneus, precuneus, and superior and inferior occipital regions, reflecting a “posterior cortical/visual neuroanatomical morphometric network” **(Fig 1**).

### Sex-Stratified Associations Between ADI and Neuroanatomical Morphometric Factor Scores

Linear regression results for male and female participants are presented in **Tables 2** and **3**, respectively. In male patients, greater neighborhood-level socioeconomic disadvantage (higher ADI national percentile rank) was significantly associated with lower neuroanatomical morphometric factor scores for three of the five neuroanatomical morphometric networks after FDR correction. Specifically, higher ADI was associated with lower cerebellar factor scores (ML1: β = −0.006, 95% CI [−0.009, −0.003], *P* < .001), lower frontal/executive factor scores (ML2: β = −0.004, 95% CI [−0.007, −0.0004], *P* = .04), and lower medial temporal/limbic factor scores (ML4: β = −0.004, 95% CI [−0.007, −0.001], *P* = .01; **Table 2**). No significant associations were observed for the subcortical-ventricular (ML3: *P* = .63) or posterior cortical/visual (ML5: *P* = .81) factors (**Table 2**). In contrast, no significant associations between ADI national percentile rank and any neuroanatomical morphometric factor score were observed in female participants after FDR correction (all *P*s ≥ .62; **Table 3**).

**Table 2:** Linear regression results: ADI national percentile rank and neuroanatomical morphometric factor scores in Male Patients (*n* = 1,112)

| Factor / Predictor | β-value | SE | 95% CI | P-value |
| --- | --- | --- | --- | --- |
| **ML1: Cerebellar** |  |  |  |  |
| ADI national percentile | -0.006 | 0.002 | [-0.009, -0.003] | < .001 |
| Age, years | -0.012 | 0.002 | [-0.015, -0.008] | < .001 |
| **ML2: Frontal/Executive** |  |  |  |  |
| ADI national percentile | -0.004 | 0.002 | [-0.007, -0.0004] | .04 |
| Age, years | -0.015 | 0.002 | [-0.018, -0.012] | < .001 |
| **ML3: Subcortical-Ventricular** |  |  |  |  |
| ADI national percentile | -0.001 | 0.002 | [-0.004, 0.002] | .63 |
| Age, years | -0.009 | 0.002 | [-0.012, -0.006] | < .001 |
| **ML4: Medial Temporal/Limbic** |  |  |  |  |
| ADI national percentile | -0.004 | 0.001 | [-0.007, -0.001] | .01 |
| Age, years | -0.018 | 0.001 | [-0.021, -0.015] | < .001 |
| **ML5: Posterior Cortical/Visual** |  |  |  |  |
| ADI national percentile | -0.0004 | 0.002 | [-0.003, 0.003] | .81 |
| Age, years | 0.003 | 0.002 | [0.0003, 0.006] | .03 |

**Note:**— $\beta$ = unstandardized regression coefficient; SE = standard error; CI = confidence interval; ADI = area deprivation index; ML = maximum likelihood factor; FDR = false discovery rate. FDR correction was applied across the five ADI coefficient *P*-values within each sex using the Benjamini-Hochberg method. Age was included as a covariate in all models.

**Table 3:** Linear regression results: ADI national percentile rank and neuroanatomical morphometric factor scores in female patients (*n* = 1,751)

| Factor / Predictor | β-value | SE | 95% CI | P-value |
| --- | --- | --- | --- | --- |
| **ML1: Cerebellar** |  |  |  |  |
| ADI national percentile | -0.001 | 0.001 | [-0.004, 0.001] | .62 |
| Age, years | -0.010 | 0.001 | [-0.013, -0.008] | < .001 |
| **ML2: Frontal/Executive** |  |  |  |  |
| ADI national percentile | -0.002 | 0.001 | [-0.004, 0.001] | .62 |
| Age, years | -0.008 | 0.001 | [-0.010, -0.006] | < .001 |
| **ML3: Subcortical-Ventricular** |  |  |  |  |
| ADI national percentile | 0.001 | 0.001 | [-0.002, 0.003] | .62 |
| Age, years | -0.004 | 0.001 | [-0.006, -0.001] | .002 |
| **ML4: Medial Temporal/Limbic** |  |  |  |  |
| ADI national percentile | -0.001 | 0.001 | [-0.003, 0.002] | .62 |
| Age, years | -0.015 | 0.001 | [-0.018, -0.013] | < .001 |
| **ML5: Posterior Cortical/Visual** |  |  |  |  |
| ADI national percentile | -0.001 | 0.001 | [-0.004, +0.001] | .62 |
| Age, years | 0.006 | 0.001 | [0.003, 0.008] | < .001 |

**Note:**— $\beta$ = unstandardized regression coefficient; SE = standard error; CI = confidence interval; ADI = area deprivation index; ML = maximum likelihood factor; FDR = false discovery rate. FDR correction was applied across the five ADI coefficient *P*-values within each sex using the Benjamini-Hochberg method. Age was included as a covariate in all models.

## DISCUSSION

In this retrospective cross-sectional neuroimaging study of 2,863 real-world clinical patients, we investigated whether neighborhood-level socioeconomic disadvantage was associated with brain morphometry in a sex-stratified manner. Using exploratory factor analysis to reduce 131 intracranial volume-normalized regional brain volumes into five latent neuroanatomical morphometric networks, and subsequent sex-stratified linear regression, we identified associations between ADI national percentile rank and neuroanatomical morphometric factor scores in male but not female patients. Specifically, greater neighborhood disadvantage was associated with lower factor scores across cerebellar, frontal/executive, and medial temporal/limbic networks in male patients, with no significant associations observed in female patients (**Tables 2** and **3**). These findings extend prior work from our group^11^ by revealing that the neuroanatomical morphometric correlates of neighborhood disadvantage may be sex-specific and span multiple distributed neuroanatomical morphometric networks detectable in routine clinical neuroimaging.

The observation that ADI was significantly associated with regional morphometry in male but not female patients is a notable finding. This is consistent with recent evidence suggesting that the neurobiological embedding of socioeconomic disadvantage is not uniform across sexes. Prior work in disease-enriched samples have begun to demonstrate sex-specific associations between neighborhood disadvantage and cortical morphometry, including regions implicated in stress reactivity, emotion regulation, and pain processing.^6,10^ The present findings extend this literature to a broader, real-world clinical population, and demonstrate that sex-specificity is observable at the level of latent neuroanatomical morphometric networks rather than individual regions.

Several mechanisms may account for the differential susceptibility observed in male patients. Biological sex modulates stress response physiology across the lifespan, including hypothalamic-pituitary-adrenal axis reactivity, glucocorticoid sensitivity, and overall autonomic nervous system function, all of which have downstream effects on brain structure.^13^ Males and females also differ in the nature, timing, and neurobiological consequences of the social exposure, and these differences may render male brain morphometry more sensitive to the chronic, ambient stressors associated with neighborhood disadvantage.^12,13^ Furthermore, sex differences in cardiovascular risk trajectories,^15^ which are in and of themselves potentiated by neighborhood disadvantage,^6,7,11^ may contribute to differential patterns of morphometric vulnerability, particularly in metabolically active brain regions including the prefrontal cortex and cerebellum.^30–32^

Among the three factors significantly associated with ADI in male participants, the cerebellar neuroanatomical morphometric network (ML1) demonstrated the strongest association. This finding is noteworthy given that the cerebellum is increasingly recognized as a contributor to cognitive, affective, and motor regulation extending beyond its classical sensorimotor role,^33,34^ is sensitive to both early life adversity and the life-course social exposure.^35,36^ To our knowledge, no study to date has identified associations between ADI and cerebellar morphometry.^5–11^ The medial temporal/limbic neuroanatomical morphometric network, encompassing the hippocampus, amygdala, entorhinal cortex, and parahippocampal gyrus, contains structures with well-established roles in memory consolidation, emotional processing, and stress regulation,^37,38^ and which have been among the most consistently identified targets of socioeconomic adversity in prior neuroimaging research.^5,6,8,39^ The frontal/executive neuroanatomical morphometric network, anchored by dorsolateral prefrontal cortex and extending across middle and superior frontal gyri, supplementary motor cortex, and orbitofrontal cortex; regions with numerous high-level functions including working memory, cognitive control, and behavioral regulation.^40–43^ Reduced frontal morphometry in the context of greater neighborhood disadvantage is consistent with a broader literature linking socioeconomic adversity to prefrontal structural and functional compromise.^5,44,45^ Conversely, the subcortical-ventricular (ML3) and posterior cortical/visual (ML5) neuroanatomical morphometric networks were not significantly associated with ADI in either sex, suggesting that the neuroanatomical correlates of neighborhood disadvantage may be, to an extent, preferentially expressed in phylogenetically newer and/or developmentally protracted brain regions.^46–49^

Our findings build directly on our prior results demonstrating that ADI was associated with higher brain age gap, lower total brain tissue volume, and amplified associations between white matter hyperintensity burden and regional volumes in this clinical sample.^11^ The present study extends this work in two key respects: first, by disaggregating whole-brain morphometric associations into network-level morphometric patterns using factor analysis, and second, by demonstrating that these ADI–morphometric associations differ by sex. The absence of any main effects of ADI on regional morphometry in prior work, where sex was included as a covariate, but analyses were not stratified, is consistent with the null result observed in females in the present study potentially diluting sex-pooled effects.^11^ The present sex-stratified design was specifically motivated by this possibility, and the results support this vein of analysis.

These findings should be considered in the context of several limitations. First, this study is cross-sectional, precluding causal inference or assessing cumulative disadvantage exposure over the life course. Second, the retrospective epidemiological structure of the study increases the potential for residual confounding, given that there were numerous demographic variables that we were unable to account for. Third, the cohort is geographically restricted to a single academic medical center and its community partners in a single state, which may limit generalizability to other regions of the country or health system contexts within the state. Fourth, the residential addresses at the time of the clinical encounter may also not accurately reflect lifetime neighborhood context, and individuals who have recently relocated may be misclassified with respect to chronic exposure, which may be particularly consequential for brain development and morphometry. Finally, sex was operationalized here as binary biological sex recorded from the medical record. Gender identity, and its potential interactions with the social exposome and brain health, were therefore unable to be assessed.^50^

Nevertheless, these findings carry implications for both neuroimaging research and clinical practice. From a methodological standpoint, the results reinforce the importance of sex-stratified analyses in studies of socioeconomic determinants of brain health. Pooled analyses may systematically mask sex-specific associations or produce estimates that do not generalize to either sex independently. From a clinical perspective, the identification of cerebellar, frontal/executive, and medial temporal/limbic neuroanatomical morphometric networks being differentially associated with neighborhood disadvantage in male patients may help contextualize morphometric findings in clinical neuroimaging, particularly in males from socioeconomically disadvantaged communities who present without overt structural pathology. Future work examining the longitudinal trajectories of these associations, and whether they are modifiable by neighborhood-level intervention, would further clarify the mechanisms through which structural inequities become reflected in brain morphometry across the lifespan.

## CONCLUSIONS

In a large, real-world clinical population, greater neighborhood-level socioeconomic disadvantage was associated with lower neuroanatomical morphometric factor scores across cerebellar, frontal/executive, and medial temporal/limbic neuroanatomical morphometric networks in male but not female patients, reflecting smaller regional brain volumes within these distributed networks. Taken together, these findings suggest that the neuroanatomical correlates of neighborhood disadvantage are not uniform across biological sexes, and that sex-stratified analyses may be necessary to fully characterize the relationship between the social exposome and brain morphometry in clinical populations. Altogether, integrating neighborhood-level socioeconomic context into the interpretation of clinical neuroimaging data will be critical to improve the accuracy and equity of brain health assessment across diverse patient populations.

## ABBREVIATIONS

ADI: area deprivation index
FDR: false discovery rate
IQR: interquartile range
KMO: Kaiser-Meyer-Olkin
ML: maximum likelihood factor

