## Supplemental Material for "Sex-Specific Regional Brain Morphometric Correlates of Neighborhood Socioeconomic Disadvantage in Clinical Neuroimaging"

**Table S1:**

Regions of interest included in factor analysis (*n* = 131)

**Region of Interest**

**Cerebellar** (*n* = 5)

Cerebellum Exterior (R)

Cerebellum Exterior (L)

Cerebellar Vermal Lobules I–V (total)

Cerebellar Vermal Lobules VI–VII (total)

Cerebellar Vermal Lobules VIII–X (total)

**Subcortical** (*n* = 12)

Accumbens Area (R)

Accumbens Area (L)

Caudate (R)

Caudate (L)

Pallidum (R)

Pallidum (L)

Putamen (R)

Putamen (L)

Thalamus Proper (R)

Thalamus Proper (L)

Ventral Diencephalon (R)

Ventral Diencephalon (L)

**Ventricular / Subcallosal** (*n* = 6)

Lateral Ventricle (R)

Lateral Ventricle (L)

Inferior Lateral Ventricle (R)

Inferior Lateral Ventricle (L)

Subcallosal Area (R)

Subcallosal Area (L)

**Frontal / Motor** (*n* = 40)

Anterior Orbital Gyrus (R)

Anterior Orbital Gyrus (L)

Central Operculum (R)

Central Operculum (L)

Dorsolateral Prefrontal Cortex (R)

Dorsolateral Prefrontal Cortex (L)

Frontal Operculum (R)

Frontal Operculum (L)

Frontal Pole (R)

Frontal Pole (L)

Gyrus Rectus (R)

Gyrus Rectus (L)

Lateral Orbital Gyrus (R)

Lateral Orbital Gyrus (L)

Medial Frontal Cortex (R)

Medial Frontal Cortex (L)

Medial Orbital Gyrus (R)

Medial Orbital Gyrus (L)

Middle Frontal Gyrus (R)

Middle Frontal Gyrus (L)

Opercular Part of Inferior Frontal Gyrus (R)

Opercular Part of Inferior Frontal Gyrus (L)

Orbital Part of Inferior Frontal Gyrus (R)

Orbital Part of Inferior Frontal Gyrus (L)

Orbitofrontal Cortex (R)

Orbitofrontal Cortex (L)

Posterior Orbital Gyrus (R)

Posterior Orbital Gyrus (L)

Precentral Gyrus (R)

Precentral Gyrus (L)

Precentral Gyrus Medial Segment (R)

Precentral Gyrus Medial Segment (L)

Superior Frontal Gyrus (R)

Superior Frontal Gyrus (L)

Superior Frontal Gyrus Medial Segment (R)

Superior Frontal Gyrus Medial Segment (L)

Supplementary Motor Cortex (R)

Supplementary Motor Cortex (L)

Triangular Part of Inferior Frontal Gyrus (R)

Triangular Part of Inferior Frontal Gyrus (L)

**Temporal / Medial Temporal / Limbic** (*n* = 30)

Amygdala (R)

Amygdala (L)

Anterolateral Temporal Cortex (R)

Anterolateral Temporal Cortex (L)

Entorhinal Area (R)

Entorhinal Area (L)

Fusiform Gyrus (R)

Fusiform Gyrus (L)

Hippocampus (R)

Hippocampus (L)

Inferior Temporal Gyrus (R)

Inferior Temporal Gyrus (L)

Mediotemporal Cortex (R)

Mediotemporal Cortex (L)

Middle Temporal Gyrus (R)

Middle Temporal Gyrus (L)

Occipital Fusiform Gyrus (R)

Occipital Fusiform Gyrus (L)

Parahippocampal Gyrus (R)

Parahippocampal Gyrus (L)

Planum Polare (R)

Planum Polare (L)

Planum Temporale (R)

Planum Temporale (L)

Superior Temporal Gyrus (R)

Superior Temporal Gyrus (L)

Temporal Pole (R)

Temporal Pole (L)

Transverse Temporal Gyrus (R)

Transverse Temporal Gyrus (L)

**Parietal** (*n* = 12)

Angular Gyrus (R)

Angular Gyrus (L)

Parietal Operculum (R)

Parietal Operculum (L)

Postcentral Gyrus (R)

Postcentral Gyrus (L)

Postcentral Gyrus Medial Segment (R)

Postcentral Gyrus Medial Segment (L)

Superior Parietal Lobule (R)

Superior Parietal Lobule (L)

Supramarginal Gyrus (R)

Supramarginal Gyrus (L)

**Occipital / Visual** (*n* = 16)

Calcarine Cortex (R)

Calcarine Cortex (L)

Cuneus (R)

Cuneus (L)

Inferior Occipital Gyrus (R)

Inferior Occipital Gyrus (L)

Lingual Gyrus (R)

Lingual Gyrus (L)

Middle Occipital Gyrus (R)

Middle Occipital Gyrus (L)

Occipital Pole (R)

Occipital Pole (L)

Precuneus (R)

Precuneus (L)

Superior Occipital Gyrus (R)

Superior Occipital Gyrus (L)

**Insular** (*n* = 4)

Anterior Insula (R)

Anterior Insula (L)

Posterior Insula (R)

Posterior Insula (L)

**Cingulate** (*n* = 6)

Anterior Cingulate Gyrus (R)

Anterior Cingulate Gyrus (L)

Middle Cingulate Gyrus (R)

Middle Cingulate Gyrus (L)

Posterior Cingulate Gyrus (R)

Posterior Cingulate Gyrus (L)

**Note:—**All 131 regions of interest are intracranial volume-normalized bilateral regional volumes derived from an automated atlas-based segmentation (cNeuro, Combinostics Oy).

(L) = left hemisphere; (R) = right hemisphere.

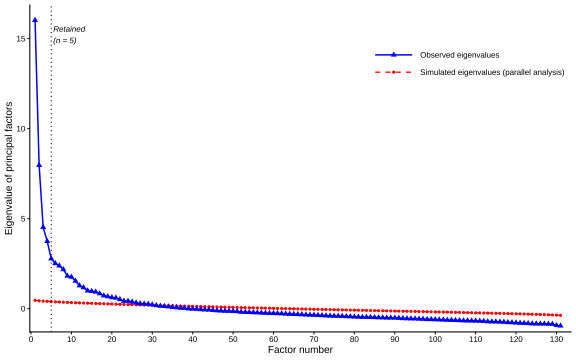

**FIG S1.**

Parallel analysis scree plot supports retention of five factors based on scree plot inflection despite a formal suggestion of 30 factors. Scree plot from parallel analysis of the 131-region brain volume correlation matrix (*n* = 2,863). Solid blue line with triangles indicates observed eigenvalues of principal factors; dashed red line indicates simulated eigenvalues derived from 100 iterations of parallel analysis using a maximum likelihood estimator. Parallel analysis formally suggested retention of 30 factors at the point where observed eigenvalues exceed simulated eigenvalues; however, a clear inflection point is observed after the fifth factor. Accordingly, five factors were retained based on scree plot inspection and anatomical interpretability of the resulting solution. Eigenvalues are plotted on the y-axis; factor number on the x-axis.

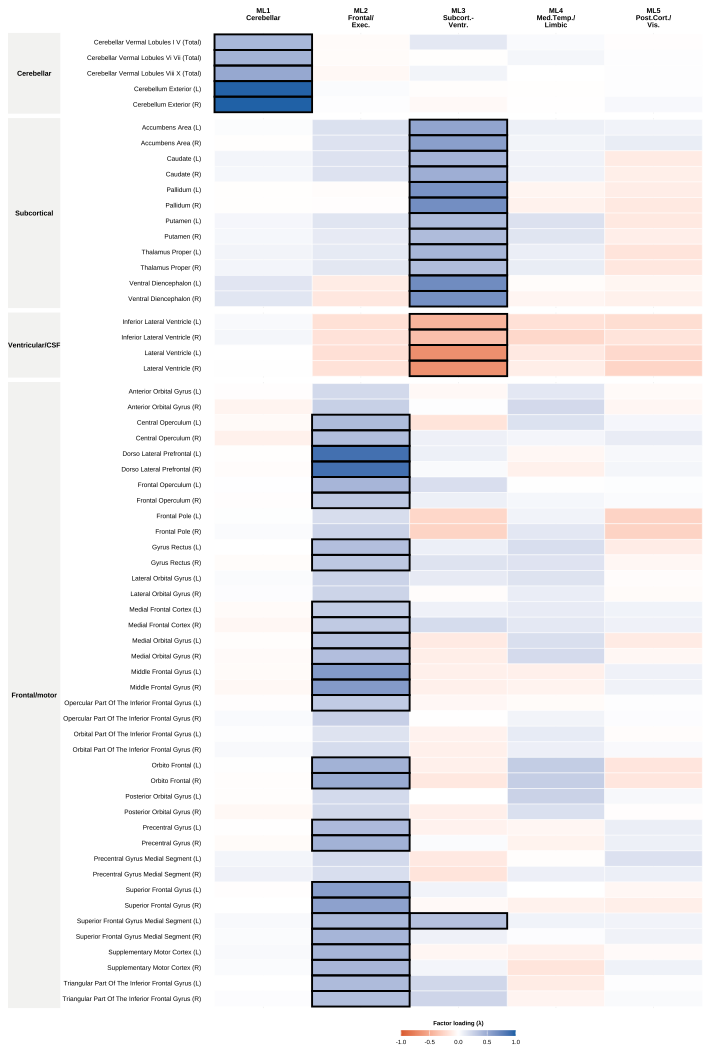

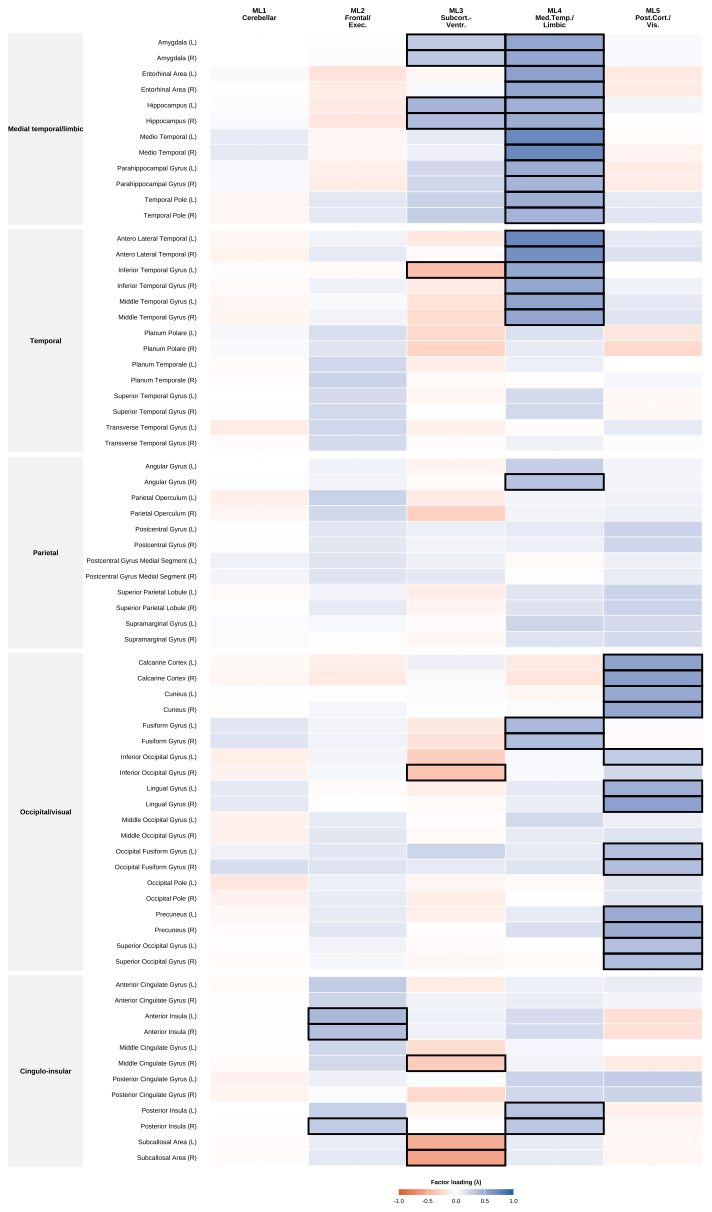

**FIG S2.**

Full factor loadings for all 131 regions of interest. All intracranial volume-normalized regional brain volumes shown, grouped by neuroanatomical category (as depicted in Table S1). Blue shading indicates positive loadings; orange shading indicates negative loadings; color intensity indicates loading magnitude. Bold cell borders indicate |λ| ≥ 0.30, consistent with established conventions for minimum factor loading interpretation.^1–3^

ML = maximum likelihood factor; λ = factor loading.
